# Efficient reverse breeding by VIGS-mediated transient crossover reduction

**DOI:** 10.1101/459016

**Authors:** Vanesa Calvo-Baltanas, Cris L. Wijnen, Nina Lukhovitskaya, C. Bastiaan de Snoo, Linus Hohenwarter, Hans de jong, Arp Schnittger, Erik Wijnker

## Abstract

F1 heterozygotes are traditionally generated by crossing homozygous parental lines. The opposite is achieved through reverse breeding, in which parental lines are generated from a heterozygote. Reverse breeding can be used to develop new F1 hybrid varieties without having prior access to homozygous breeding lines. For successful reverse breeding, the heterozygotes’ homologous chromosomes must be divided over two haploid complements, which is achieved by suppression of meiotic crossover (CO) recombination. We here show two innovations that facilitate efficient reverse breeding. Firstly, we demonstrate that downregulation of CO rates can be accomplished using virus-induced gene silencing (VIGS). We obtain transgene-free parental lines for a heterozygote in just two generations. Secondly, we show that incomplete CO suppression opens up several alternative strategies for the preservation of hybrid phenotypes through reverse breeding.

---

Heterozygous F1 hybrids are among the highest producing crop varieties^1^ and result from intercrossing homozygous parental lines. Existing hybrids are usually further improved through the introgression of new alleles into their parental lines. In an alternative approach, large numbers of new and potentially better heterozygous genotypes could be generated in outcrossing populations, for example by intercrossing different commercially available heterotic hybrids and selecting the best performing heterozygotes in their offspring. However, this potential is rarely, if ever exploited, because unique heterozygotes selected from outcrossing populations cannot be maintained: when they set seed, their unique allele combinations are lost through meiotic recombination. This restriction can be overcome by reverse breeding, in which new parental lines for any heterozygote can be *post-hoc* generated from the selected heterozygote itself^2,3^ (Fig. 1). By obtaining its parental lines, a heterozygote can be recreated as F1 hybrid.

**Figure 1.**
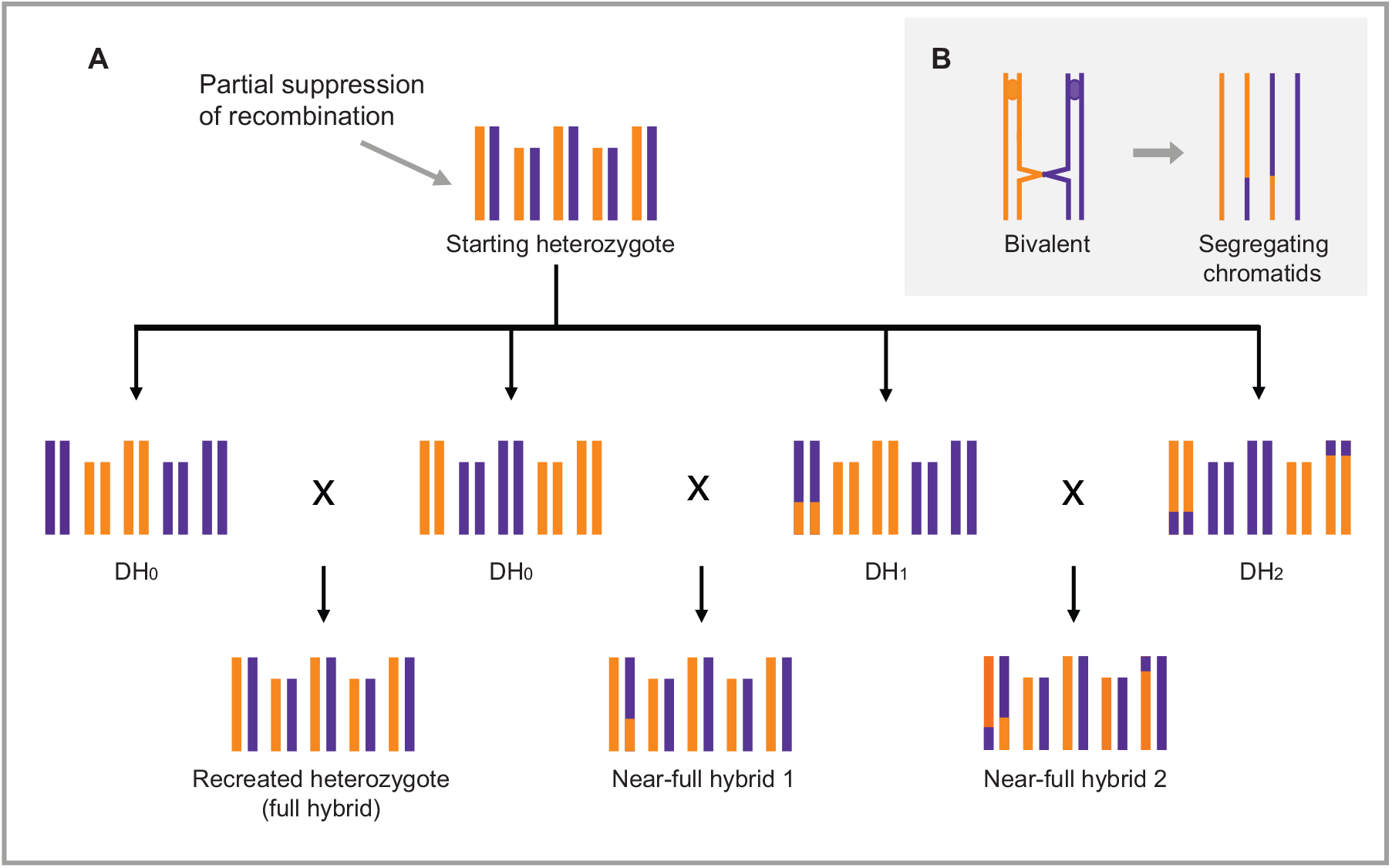
Reverse breeding (heterozygote reconstruction) and near-full hybrid generation through partial crossover (CO) suppression. Panel A: a starting Arabidopsis heterozygote (top), for which parental lines are to be made. Five chromosome pairs are shown, with homologs in orange and purple. Meiotic COs are partially suppressed in this heterozygote, resulting in DH offspring as shown in the middle row having 0, 1 or 2 COs (DH_0_, DH_1_ and DH_2_ respectively). Intercrossing complementing DH_0_ reconstitutes the heterozygote as a full hybrid (left), which is similar to the approach by Wijnker et al., 2012^3^. Intercrossing DH_0_ with DH_1_ (middle) or DH_1_ with DH_2_ (right) generates near-full hybrids, which have small homozygous genomic regions. Note for example that in the cross of DH_1_ with DH_2_ chromosome 1 is largely heterozygous, since the parental lines complement one another in the distal chromosome region. Panel B: Recombinant but also non-recombinant chromatids can segregate in the presence of CO. Detail of a bivalent pair with one meiotic CO is shown (left). Only two of the four chromatids are recombinant (right).

A proof of concept study^3^ showed the feasibility of reverse breeding in an *Arabidopsis thaliana* hybrid. This was achieved by the complete *knock-down* of meiotic CO formation in a F1 hybrid using a dominantly acting RNAi transgene targeting the essential meiotic recombinase DISRUPTED MEIOTIC CDNA 1 (DMC1). Without COs, non-recombinant chromosomes segregate to gametes. These gametes were regenerated as haploid plants, and self-fertilized to give rise to homozygous diploid lines (doubled haploids; DH) from which complementing parental lines were selected and crossed to reconstitute the starting heterozygote (Fig. 1A). In short, reverse breeding requires the consecutive suppression of recombination and the conversion of resulting gametes to DH offspring.

Since the translation of this technique to crops may be challenging, we here set out to overcome two major drawbacks of the original approach. Firstly, the use of a transgene to suppress CO formation in a heterozygote is impractical. Stable transformation of a selected heterozygote can be complex and a transgene that dominantly compromises fertility renders half of the offspring (the genotypes carrying the construct) useless for further breeding. We asked if virus-induced gene silencing (VIGS) could be used to transiently suppress meiotic CO formation in a hybrid^4–6^ and whether gametes resulting from VIGS-modified meiosis can be used to generate offspring of desired genotypic composition.

Secondly, CO formation is indispensable for chromosome segregation in plants. Without COs, homologs segregate randomly (as univalents) at anaphase I. This causes aneuploidy in gametes and semi-sterility. Viable haploid gametes can still be formed in the absence of COs, when the homologues of each chromosome pair by random chance segregate to opposite poles^3^. The probability of regular disjunction is a function of the chromosome number of the plant^2^ (n): P_(balanced segregation)_=1/2^n^. The more chromosome pairs, the lower the probability of viable gamete formation, and the lower the chance of obtaining parental lines. In Arabidopsis about 1/2^5^ = 3% of meiotic events generates viable spores in the absence of COs.

The suppression of CO formation enriches for the segregation of non-recombinant chromosomes to gametes, but complete CO suppression is not essential. Gametes carrying exclusively non-recombinant chromosomes will occasionally be formed in wild-type meiosis (see Fig. 1B) although they are usually rare, especially when the chromosome number is high (see Supplementary tables 1-6). A reduction of CO, rather than complete CO suppression, might present a favorable intermediate approach to enrich for viable gametes carrying only or mostly, non-recombinant chromosomes^2^. The presence of parallel pathways that lead to CO formation in plants allows theoretically for fine-tuning CO rates^7^. Mutants of *MUTS HOMOLOGUE5 (MSH5)* show about 87% reduction in COs in Arabidopsis^8^ and we therefore targeted *MSH5* using a VIGS construct to reduce CO formation.

The efficiency of VIGS to downregulate *MSH5* was assessed by inoculating plants at the five-leaf stage with a VIGS vector (TRV2-*AtMSH5*) and evaluating meiotic progression. *MSH5* knocked-down plants exhibited high levels of aborted pollen about three weeks after inoculation and siliques that failed to elongate (Supplementary fig. 1), consistent with a *msh5* mutant phenotype^9^. Chromosome spreads of late meiotic cell complements confirmed the mis-segregation of chromosomes during meiosis (Supplementary fig. 1). Reduced fertility was typically observed for three to four days on flowers, after which the plants reverted to a wild-type phenotype, exhibiting viable pollen and long siliques.

To evaluate the feasibility of breeding with gametes resulting from VIGS-mediated reduction of recombination, we inoculated an F1 (Landsberg *erecta* x Columbia) with TRV2-*AtMSH5*. Once the flowers showed a high fraction of aborted pollen, they were crossed to *GFP-tailswap*, a haploid inducer line for Arabidopsis^10^. Haploid offspring were obtained and self-fertilized to give rise to 111 DH offspring that were genotyped for 42 markers evenly spaced over the genome (Supplementary fig. 2).

Among the 111 offspring we identified 24 DHs (20 different genotypes) that carry only non-recombinant chromosomes (Supplementary file 1). These lines, which are also known as chromosome substitution lines (CSLs), are henceforth referred to as DH_0_. The population is significantly enriched for DH lines originating from non-recombinant chromosomes in comparison to a previously published wild-type population (Fig. 2) (Kolmogorov-Smirnov test; α=0.01). Among these 20 DH_0_ genotypes we identified six complementing parental pairs that, when crossed, recreate the starting hybrid (Supplementary file 1). All DH offspring developed normally and were fully fertile. This shows that VIGS can transiently modify meiotic recombination in wild-type hybrids. This approach strongly increases reverse breeding efficiency and speed as compared to an earlier approach in which CO suppression resulted from using a dominant RNAi transgene^3,11^. Firstly, using VIGS, all recovered offspring are transgene-free and fertile, while in the previous set-up half of the offspring were transgenic and sterile implying a two-fold increase of efficiency. Moreover, without stable transformation a heterozygous genotype can be recreated as F1 in three generations where previously six generations were required.

**Figure 2.**
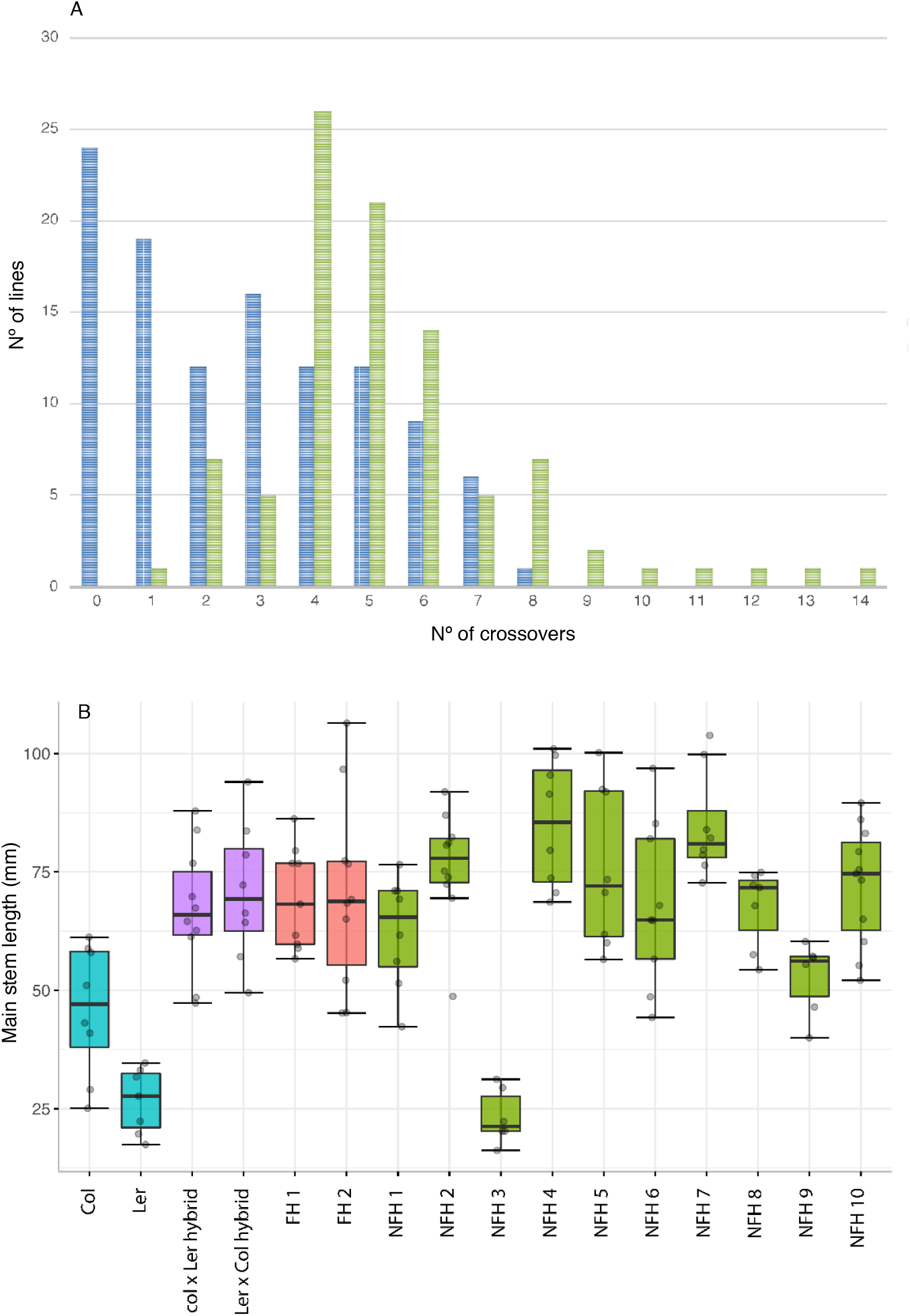
Crossover (CO) distributions in reverse breeding-and wild-type DH offspring (top panel) and comparison of hybrid phenotypes (lower panel). Top panel (A) shows the observed CO number for wild-type DH offspring (in green) and for reverse breeding DH offspring (in blue). Note that reverse breeding are enriched for DH offspring having 0 and 1 COs. Lower panel (B) shows box-and-whisker plots for main stem length at the moment of flowering of parental lines, full hybrids (FH) and near-full hybrids (NFH). Parental lines Col-0 and L*er* (shown in blue), Col-0 x L*er* reciprocal hybrids (violet), full hybrids (coral) and near-full hybrids (green). Genotypes of near full hybrids are presented in the supplementary file.

For the exact recreation of a heterozygous genotype, complementing DH_0_ are required, but in practice the recreation of the hybrid *phenotype* will be the ultimate goal. The use of a DH_1_ (i.e. a DH with one recombinant chromosome) in a cross to recreate a hybrid, leads to a decrease of heterozygosity (hereafter DOH) in the reconstituted hybrid distal to the CO position (Fig. 1). We hypothesized that only in case DOH negatively affects the hybrid phenotype, it is of concern for reverse breeding. In our offspring we identified 19 DH_1_ and 12 DH_2_ with one and two COs per genome respectively, with the remainder of 58 DH having three to eight COs, which is in the range of wild-type meiosis and likely result from incomplete penetrance of VIGS (Supplementary file 1). Depending on CO positions in the DH_1_ and DH_2_ offspring, we noted the possibility of identifying four additional parental pairs in which a near-full hybrid would show a DOH less than 2.5% of the total genome length. Seven parental pairs would give rise to near-full hybrids in which DOH is less than 5%. Only near-full hybrids with one CO show less than 2.5 % DOH. In one parental pair (DH_1_ line 44 x DH_2_ line 41) COs on the same chromosome arm partly compensate, similar to the DH_1_ x DH_2_ cross illustrated in Fig. 1, generating a near-full hybrid with a DOH of 4.2%.

The phenotypic impact of DOH can be explored experimentally. We therefore intercrossed DH lines to create near-full hybrids with increasing levels of DOH ranging from 1.28% −32.1% (Supplementary fig. 3). These were grown together with the starting heterozygote and full hybrids (recreated heterozygotes) and compared standard growth parameters: flowering time, main stem length, rosette diameter and dry weight at flowering time. No significant differences were found between the starting hybrid and the full hybrids (one-way ANOVA; FT p-value = 0.3015; MSL p-value = 0.9347; RD p-value = 0.8655; DW p-value = 0.2697; Fig. 2B; Supplementary fig. 3). Also, no significant differences between the full hybrid and the near-full hybrids were found, with the exception of one: a near-full hybrid that has a similar short stem length as one of its parental lines (Fig. 2), which is likely caused by homozygosity of the main effect *erecta* locus that is homozygous in this specific hybrid^12^.

These results mainly illustrate that DOH not necessarily negatively impacts hybrid performance. It is possible to estimate the expected DOH in near-full hybrids resulting from a single CO. Arabidopsis has five linkage groups (chromosomes). One CO recombines one linkage group (1/5^th^) and this CO exchanges anything between zero and half of the linkage group, which averages at 1/4^th^ of the linkage group (typically half a chromosome arm). Expected DOH caused by a single CO thus equals on average (1/4*1/5=) 5% of the total linkage map length. Of the ten near-full hybrids (with one CO) that we can produce, five have a DOH less than 5% in Mbp, exactly as predicted as the Arabidopsis genetic map correlates well with the physical chromosome length. The more chromosomes a species has, the lower the relative DOH resulting from one CO. In a species with ten chromosome pairs (e.g. maize) one CO causes a DOH of 2.5%. This decreases even further when, as in many species, COs locate relatively distal on chromosomes. Under such a scenario, not only DH_0_, but also DH_1_ and DH_2_ may prove worthy parental lines, provided that resulting near-full hybrids are phenotyped to assess their performance.

Our experiments show that CO formation during meiosis can be adjusted to favorable levels, by targeting *MSH5* rather than *DMC1* as was previously done. The lower the CO number, the more DH_0_ and DH_1_ occur in the offspring (Fig. 2, Supplementary tables 1-6), but also the higher the level of gamete abortion. Depending on ones’ interest in obtaining DH_0_ (and DH_1_), the optimal CO rate can be calculated to balance one against the other. Especially in species with higher chromosome numbers, such considerations matter. In a species with ten chromosome pairs, in which a typical bivalent has two COs, complete CO suppression generates 100% DH_0_ offspring but results in just 0.10% of spore viability. Reducing COs by 75% -from 20 to five COs per meiosis- would increase spore viability 32 fold to 3%, since then only five rather than ten univalent pairs segregate. Of those offspring, 5.6% are DH_0_ (supplementary table 3). This is low in comparison to complete CO suppression, but it is a substantial 60.000 fold increase in comparison to wild-type meiosis. Likewise, chances for obtaining DH_1_ in its offspring are 18.8%, equal to about 10.000 fold increase. Supplementary tables 1-6 give expected DH_0_ and DH_1_ numbers at different levels of CO suppression and different chromosome numbers. Such calculations will help to determine the best possible approach for other species than Arabidopsis.

Apart from generating parental lines for heterozygotes, reverse breeding provides a way to generate populations of DH_0_, also known as chromosome substitution lines^3^. Due to the low number of segregating loci (chromosomes), such populations are near unparalleled tools to identify QTL and map complex epistatic interactions (Wijnen *et al.*, submitted)^13^. Since in mixed DH_0_/DH_1_ populations the number of segregating loci remains near minimal, the detection power of QTLs and epistasis is unlikely to decrease much. Efficient reverse breeding strategies are therefore a way towards detection and mapping complex interactions. VIGS vectors are available for a multitude of crops^14^ and exploring these for the modification of meiosis may advance breeding strategies in other species. The recent identification of meiotic mutants may present further attractive targets for VIGS-mediated breeding strategies.

## Acknowledgements

This research was supported by the Netherlands the Organization for Scientific Research (NWO) through number STW-14389 (E.W.) and the European Community (EC) though the Marie-Curie Initial Training Network “COMREC”, project 606956 funded under FP7-PEOPLE (V.C.-B.). We thank Cilia Lelivelt (Rijk Zwaan, Fijnaart, Netherlands) for her support in processing genotyping samples and Bas Zwaan (Wageningen University, Netherlands) for moments of reflection during our research. We thank Laurens Deurhof (Wageningen University, Netherlands) and Shinichiro Komaki (NAIST, Japan) for their support and help during the experimental set-up.

## Author contributions

E.W. conceptualized the research; E.W., A.S. and H.D.J. were involved in supervision and funding acquisition. V.C.B. and E.W. planned research, performed crosses. N.L. helped with setting up VIGS experiments and construct design; Cloning and VIGS experiments were done by V.C.B.; L.H. performed cytogenetic analyses with help of E.W.; C.B.d.S. performed genotyping; V.C.B., E.W. and C.L.W. designed, performed and analyzed the phenotyping experiment. V.C.B. and E.W. processed and interpreted experimental data, designed the figures and drafted the manuscript with the help of A.S. and H.d.J. All authors discussed the results and commented on the manuscript.

## Online Methods

Plant material and growth. *Arabidopsis thaliana* plants used in crosses and for VIGS inoculation were grown in potting soil in growth chambers (Percival), 21°/18° C, 16H / 8H light cycle and 60%-50% relative humidity. Haploid offspring were grown under similar conditions in a greenhouse. For phenotyping, seeds of DH offspring, reconstituted full hybrids and near-full hybrids were vernalized by sowing on wet filter paper and placed them for several days in the dark at 4°C for four days to ensure uniform germination. Plants were grown on 4×4 cm Rockwool blocks and watered with a flooding system with a Hyponex nutrient solution three times per week in a randomized block design with five blocks and two replicates per genotype in each block. Climate chamber conditions were set to 16h/8h and 20/18°C day/night cycle, light was set to 125 μmolm-2s-1 and there was 70% relative humidity.

Plasmid construction and *Agrobacterium* inoculation. Two *MSH5* cDNA regions were amplified using primers to which *BamH*I (forward) and *Xba*I (reverse) restriction sites were added. The MSH5_F1/R1 and MSH5_F2/R2 primer pairs give fragments of 242 bp and 254 bp respectively, and were used to generate the TRV2-*AtMSH5* and TRV2*AtMSH5*_2 constructs. Both PCR products were introduced individually into the vector TRV2 (pYL156)^15^ following a classical digestion-ligation reaction. After sequence verification, the TRV2*-AtMSH5* and TRV2*-AtMSH5_*2 vectors were transformed into *Agrobacterium tumefaciens GV3101* (*pMP90*) strain. The incubation and inoculation protocol was executed as described in Nimchuk *et al*., 2000^16^. Plant *agro*inoculation was done by leaf-infiltration^17^ of TRV2*-AtMSH5* in combination with TRV1 (pYL192)^15^ or TRV2*-AtPDS* in combination with pTRV1 in a 1:1 ratio. TRV2*-AtMSH5* and TRV2*AtMSH5*_2 induced similar pollen phenotypes in inoculated plants, after which only TRV2*-AtMSH5* was used for further experiments.

Primers used:

MSH5_F1 5’-CAGGATCCAAGCCATCGATCATTTACGC -3’

MSH5_R1 5’-CATCTAGAACTTGGACTTCACTGCCCAC -3’

MSH5_F2 5’-CAGGATCCAAGCCATCGATCATTTACGC-3’

MSH5_R2 5’-CATCTAGAACTTGGACTTCACTGCCCAC -3’

Selection of TRV-*AtMSH5 agro*inoculated plants for pollen phenotyping. A total of 109 plants were inoculated with TRV2*-AtMSH5* in three consecutive experiments (52+42+15). Three non-inoculated plants were grown as negative controls in every batch as well as three to four plants in each batch that were inoculated with TRV2-*AtPDS* to silence *pds* as positive control^5^. To evaluate a successful knock-down of *MSH5*, we assessed pollen viability in flowers that opened three weeks post-inoculation and the two consecutive weeks. One anther was removed from each flower and placed on a slide with a drop of a modified Alexander stain^18^ to observe pollen viability. Pollen on control plants remained viable throughout the test periods. The number of affected flowers was not consistent. Within an inflorescence, flowers with high levels of pollen abortion usually appear consecutively, and a semi-sterile phenotype was present for about six consecutive days after the first sterile flowers appeared.

*DH production. To produce doubled haploids, F1 hybrid plants of Ler* x Col-0 plants were inoculated with TRV2-*AtMSH5* as described above. Once flowers appeared, pollen of flowers displaying high levels of dead pollen were crossed to the inducer line *GFP*tailswap^10^. Of the three consecutively grown batches 27, 19 and 15 plants were used. Other plants did not show a semi-sterile phenotype. From these plants we used 132, 77 and 60 flowers for pollination of *GFP-tailswap*. Haploid selection was done as described in Wijnker *et al*., 2014^11^. Among the 369 offspring we identified 113 haploid offspring. For 111 of these we obtained DH seeds.

Phenotypical analysis of (near-)full hybrids. At the moment of flowering, flowering time (FT) was recorded and main stem length (MSL), rosette diameter (RD) and dry weight (DW) were measured for each plant. Phenotypic data was corrected for spatial trends and block effects with the SpATS R package, and the resulting spatial corrected raw data was used for further analyses. To establish whether the intercrosses of the DH_0_ resulted in different full hybrids, these were compared with the parental wild-type F1 using one-way ANOVA. To assess the performance of the NFH in comparison with the FH, a Dunnett test was conducted in which line FH2 was used as a control line.

Cytology. F1 hybrid flower buds were sampled 18 days post-inoculation. The inflorescences were incubated in Carnoy: a 3:1 mix of glacial acetic acid (HAc) and 99,8% EtOH and kept overnight at 4 °C. inflorescences were then washed twice with 70% EtOH (in water) and stored at 4° C. Meiotic chromosome spreads were made as previously described in Ross *et al*. (1996)^19^, stained with DAPI and analyzed using a Zeiss microscope equipped with epifluorescence optics.

Calculations of expected frequencies of DH_0_ and DH_1_. To calculate the expected number of DH_0_ and DH_1_ in Supplementary tables 1-6, the expected number of non-recombinant and recombinant chromatids was first determined for one chromosome (i.e. the case in which the haploid chromosome number equals 1). If α is the number of COs per bivalent, then the chance of recovering a non-recombinant chromatid in a spore, and hence the chance of recovering a DH_0_, equals P_(DH0)_=(1/2)^α^. The chance of finding a chromatid with one CO (and hence recovering a DH_1_) equals P_(DH1)_=α(1/2)^α^. For higher haploid chromosome numbers (n), the expected number of non-recombinant chromatids equal P_(DH0)_=(1/2)^αn^ and P_(DH1)_= αn(1/2)^αn^ For CO numbers 1, P_(DH0)_=1-α/2 and P_(DH1)_=α/2. For higher haploid chromosome numbers P_(DH0)_=(1-α/2)^n^ and P_(DH1)_=nα/2(1-α/2)^(n-1)^.

